# A splice donor variant in *CCDC189* is associated with asthenospermia in Nordic Red dairy cattle

**DOI:** 10.1101/472282

**Authors:** Terhi Iso-Touru, Christine Wurmser, Heli Venhoranta, Maya Hiltpold, Tujia Savolainen, Anu Sironen, Konrad Fischer, Krzysztof Flisikowski, Ruedi Fries, Alejandro Vicente-Carrillo, Manuel Alvarez-Rodriguez, Szabolcs Nagy, Mervi Mutikainen, Jaana Peippo, Juhani Taponen, Goutam Sahana, Bernt Guldbrandtsen, Henri Simonen, Heriberto Rodriguez-Martinez, Magnus Andersson, Hubert Pausch

## Abstract

**Background:** Cattle populations are highly amenable to the genetic mapping of male reproductive traits because longitudinal data on ejaculate quality and dense microarray-derived genotypes are available for many artificial insemination bulls. Two young Nordic Red bulls delivered sperm with low progressive motility (i.e., asthenospermia) during a semen collection period of more than four months. The bulls were related through a common ancestor on both their paternal and maternal ancestry. Thus, a recessive mode of inheritance of asthenospermia was suspected.

**Results:** Both bulls were genotyped at 54,001 SNPs using the Illumina BovineSNP50 Bead chip. A scan for autozygosity revealed that they were identical by descent for a 2.98 Mb segment located on bovine chromosome 25. This haplotype was not found in the homozygous state in 8,557 fertile bulls although five homozygous haplotype carriers were expected (P=0.018). Whole genome-sequencing uncovered that both asthenospermic bulls were homozygous for a mutation that disrupts a canonical 5’ splice donor site of *CCDC189* encoding the coiled-coil domain containing protein 189. Transcription analysis showed that the derived allele activates a cryptic splice site resulting in a frameshift and premature termination of translation. The mutated CCDC189 protein is truncated by more than 40%, thus lacking the flagellar C1a complex subunit C1a-32 that is supposed to modulate the physiological movement of the sperm flagella. The mutant allele occurs at a frequency of 2.5% in Nordic Red cattle.

**Conclusions:** Our study in cattle uncovered that CCDC189 is required for physiological movement of sperm flagella thus enabling active progression of spermatozoa and fertilization. A direct gene test may be implemented to monitor the asthenospermia-associated allele and prevent the birth of homozygous bulls that are infertile. Our results have been integrated in the Online Mendelian Inheritance in Animals (OMIA) database (https://omia.org/OMIA002167/9913/).

## Background

Impaired male fertility is a well-recognized and highly heterogeneous condition in many species including humans, mice and cattle (*e.g.*, [1],[2],[3],[4],[5], OMIA 000562-9913). Males with reduced reproductive ability may produce ejaculates with anomalous characteristics such as low sperm count, impaired progressive motility or morphological aberrations of the spermatozoa [6]. However, standard semen parameters are only poor predictors of male reproductive ability because many fertility disorders are caused by aberrations of the spermatozoa that are not immediately apparent or regularly monitored (e.g., [7],[8],[4]).

The normal movement of the sperm flagellum, i.e., activated and hyperactivated motility, facilitates active progression of spermatozoa through the oviduct. Morphological aberrations of the sperm tail such as stump tail, short tail, dysplasia of the fibrous sheath or disorganized flagellar microtubules may compromise sperm motility and fertilization success [9]. Genetic variants causing morphological abnormalities of the sperm tail have been identified in several species including humans [10],[11], pigs [12], cattle [13], and mice [14] where they impede motility. The fertilization success of ejaculates containing a high proportion of spermatozoa with low progressive motility (i.e., asthenospermia) is often reduced, both *in vivo* and *in vitro* [15],[16], albeit fertilization anecdotally being reported as possible *in vivo* [17].

Due to the low heritability of male reproductive performance, detecting genetic variants that underpin impaired male fertility requires large cohorts with phenotypic records that, moreover, have been genotyped [18]. Comprehensive data on male fertility have become available for many cattle populations due to the wide-spread use of artificial insemination (AI) [19]. The availability of both longitudinal data on semen quality and dense microarray-derived genotypes makes cattle populations highly amenable to the mapping of genetic determinants that underpin male reproductive performance [4],[13],[20],[21].

Prospective bulls for AI are selected at an early age and reared at AI centers under controlled conditions. When the bulls reach an age of approximately one year, their semen is collected once or twice per week. Immediately after semen collection, macroscopic, microscopic and computer-assisted analyses are carried out to assess whether the ejaculates are suitable for AI [22]. Up to 20% of all collected ejaculates are rejected because they do not fulfill the requirements for AI [23]. The most common pathological findings in ejaculates of insufficient quality include absence or low number of spermatozoa, reduced sperm motility, impaired sperm viability or aberrant morphology of heads and/or tails in many spermatozoa [24]. Usually, it takes a few weeks until young bulls produce semen that is considered suitable for AI, owing to individual differences in epididymal sperm maturation [22]. Young bulls that permanently produce ejaculates that do not fulfill the requirements for AI are sporadically noticed at semen collection centers. It is plausible that some of these bulls carry deleterious genetic variants that are responsible for the low semen quality they deliver (e.g., [4],[13]).

Here we present the phenotypic and genetic analysis of a recessively inherited form of sterility in Nordic Red bulls, which primarily manifests itself through asthenospermia. The availability of dense genotype data of two affected bulls enabled us to localize the disorder within a 2.98 Mb interval on bovine chromosome 25. Whole-genome sequencing of affected bulls revealed that a mutation disrupting a canonical splice donor site of the *CCDC189* gene is the most likely cause of the observed condition.

## Results

### Aberrant mitochondrial activity in ejaculates of asthenospermic bulls

Two young bulls from the Nordic Red dairy cattle breed producing ejaculates with immotile spermatozoa were noticed at a semen collection center. The average ejaculate volumes (3.16 ml) and sperm concentrations (1.94 Mio/µl) were within physiological ranges for bulls of their age. More than 80% of the spermatozoa in the ejaculates of both bulls were classified as viable and had typical morphology. However, less than 5% of the spermatozoa were progressively motile, thus not fulfilling the requirements for AI, whose threshold is at least 40%. Apart from producing ejaculates of insufficient quality, both young bulls were healthy and showed no apparent signs of disease.

Frozen semen samples from both asthenospermic bulls were thawed and subjected to computer-assisted motility analysis and flow cytometric studies. Post-thawing, the velocity of the spermatozoa was between 10 and 14 µm/s. The velocity of spermatozoa in normospermic control bulls was between 38 and 48 µm/s (sperm with progressive motility typically move at a velocity faster than 25 µm/s). In two normospermic control bulls, more than 50% of the spermatozoa showed progressive motility after thawing while in the affected bulls, the proportion of motile spermatozoa was less than 10%. The discrepancy between computer-assisted and subjective motility evaluations (10 vs 5%) results from spermatozoa displaying minor movement, i.e., the sperm tails bended peculiarly which was not classified as progressive motility in subjective microscopic sperm evaluation (**Figure 1**, **Additional file 3 & 4**). The ATP content was numerically higher, though not significant, in semen of asthenospermic than normospermic bulls. As expected, flow cytometry analysis revealed that viable post-thawed spermatozoa examined in either bull type had a tendency for destabilizing changes in the plasmalemma. However, the asthenospermic bulls had more spermatozoa with high mitochondrial membrane potential indicating increased mitochondrial activity. This leads to more spermatozoa producing superoxide and reactive oxygen species. This in turn causes a higher proportion of spermatozoa with lipid peroxidation in asthenospermic than normospermic bulls (**Table 1**). Transmission electron microscopy of the sperm flagella from asthenospermic bulls showed the typical arrangement of nine outer microtubule doublets surrounding the central pair (9×2 + 2). No ultrastructural abnormalities of the axonemes were immediately apparent (**Figure 1**).

**Table 1.** Results of the flow cytometric analysis of frozen thawed semen samples.

| Animal | Early membrane destabilization | Increased mitochondrial activity | Increased mitochondrial superoxide production | Increased ROS production | Lipid peroxidation degree |
| --- | --- | --- | --- | --- | --- |
| Asthenospermic-1 | 2.0 | 72.3 | 24.9 | 39.0 | 22.0 |
| Asthenospermic-2 | 4.2 | 71.9 | 25.9 | 17.8 | 23.3 |
| Normospermic-1 | 10.7 | 47.3 | 4.9 | 5.1 | 3.1 |
| Normospermic-2 | 8.3 | 54.3 | 1.8 | 6.7 | 3.2 |

Proportions (%) of viable spermatozoa post-thawing from two asthenospermic and two normospermic bulls of the Nordic Red breed depicting sperm early membrane destabilization (YO-PRO-1+/PI-), degree of mitochondrial activity (Mitotracker+/PI-), and increased production of mitochondrial superoxide (MitoSOX+/PI-), reactive oxygen species (ROS, CellROX+) and lipid peroxidation (Bodipy-C11 + (green)) assessed using flow cytometry.

**Figure 1.**
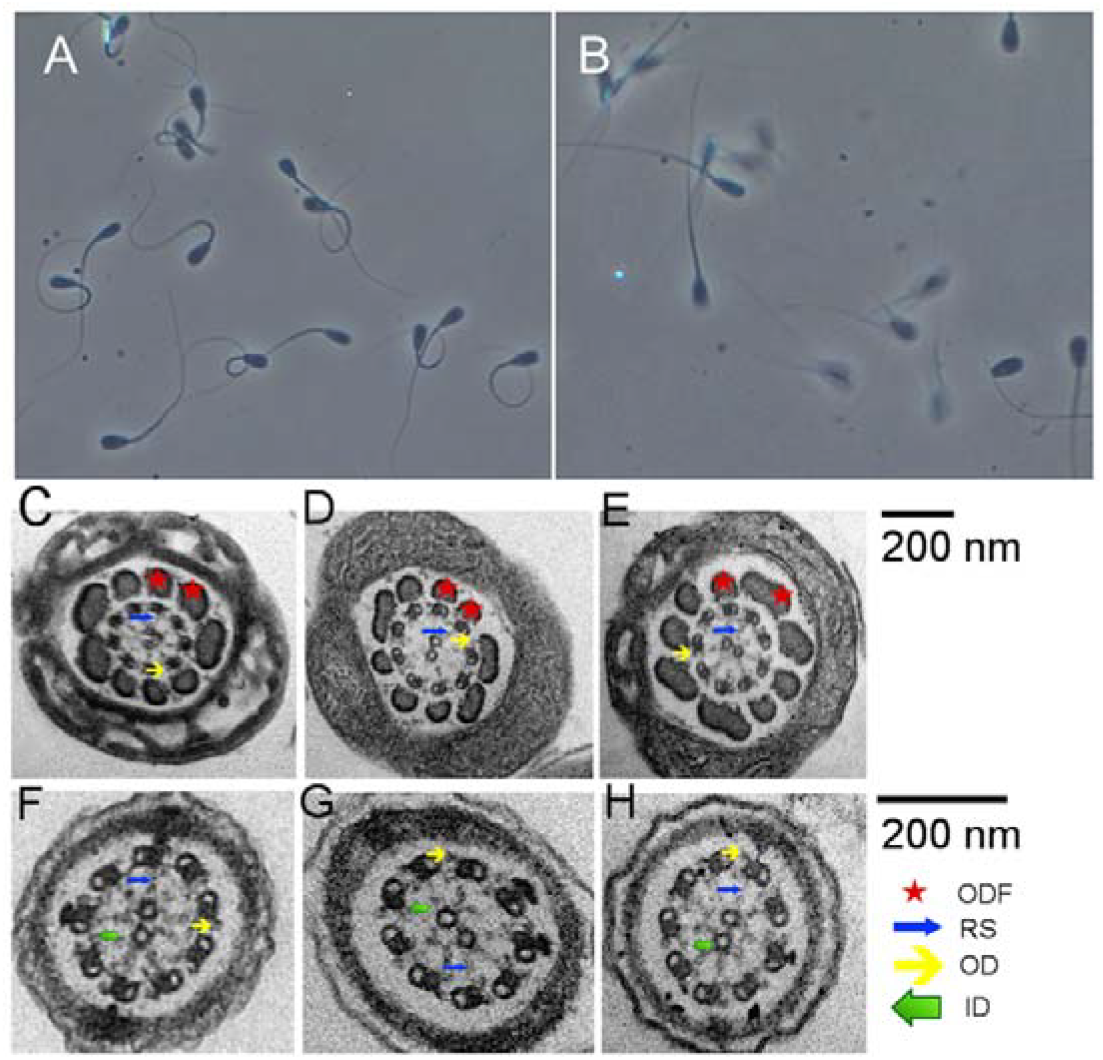
Phenotypic manifestation of asthenospermia. Microscopic view of a representative sample of frozen-thawed spermatozoa from an asthenospermic (A) and normospermic (b) bull. Bending of many spermatozoa of the asthenospermic bull and the lack of motility projected as clear pictures of spermatozoa (A). Foggy (double pictured) representation of many spermatozoa in the normospermic bull indicate actively moving spermatozoa (B). Both figures were photographed using the same settings. Transmission electron microscopy of spermatozoa from two asthenospermic (C, D, F, G) and an normospermic (E, H) bull. Representative cross-sections of the midpiece (C-E) and principal piece (F-H) of the flagella revealed the typical arrangement of nine outer microtubule doublets surrounding the central pair (9×2 + 2) in asthenospermic and normospermic bulls. No aberrations of the axonemal ultrastructures including outer dense fiber (ODF), radial spokes (RS), inner (ID) and outer dynein arms (OD) were apparent in cross-sections of asthenospermic bulls.

### Bulls with asthenospermia are sterile *in vivo* and *in vitro*

None of the 9 cows inseminated with semen of an asthenospermic bull conceived, whereas 42 of the 89 cows inseminated with normospermic semen did.

*In vitro* fertilization success was tested using 470 offal oocytes. Using post-thaw semen from an asthenospermic bull, the embryo cleavage rate at 42 hours post-insemination was only 11% (36 out of 320) compared to 70% (105 out of 150) using spermatozoa from a normospermic control bull. At day nine post-insemination, the embryo yield after embryo culture was only 0.9% (3 out of 320) for the asthenospermic bull compared to 23% (34 out of 150) for the normospermic bull. Moreover, the three embryos obtained using semen from the asthenospermic bull reached blastocyst stage only at day 9, indicating aberrantly slow development kinetics.

Because the semen analyses revealed consistent sperm deviations and deficient fertilization capacity both *in vivo* and *in vitro* in both asthenospermic bulls, we suspected that this type of sterility may represent a genetic condition. The analysis of pedigree records revealed a common male ancestor (born in 1990) that was present either as great-great-great-grandfather or great-great-grandfather in both the maternal and paternal ancestry of both affected bulls (**Additional file 5**). The sires of the two asthenospermic bulls had more than 300 and 2,000 daughters, respectively, indicating that they were frequently used for AI. According to the AI centers consulted, their semen quality was normal. Seven and twelve paternal half-sibs of the asthenospermic bulls were used for 6,257 and 36,689 artificial inseminations, respectively. According to the AI center, their semen qualities were normal and their average calving rates (number of births per insemination) were 0.39 and 0.44, i.e., similar to average values in Nordic Red cattle. Because a common male ancestor was in the maternal and paternal ancestry of both asthenospermic bulls and fertility was normal in their first- and second-degree relatives, we suspected that the disorder might follow an autosomal recessive mode of inheritance.

### Asthenospermia maps to a 2.98 Mb segment on bovine chromosome 25

To identify the genomic region associated with asthenospermia, the two affected bulls were genotyped using the BovineSNP50 Bead chip. Following quality control, genotypes of 41,594 autosomal SNPs were available for genetic analyses. Assuming recessive inheritance of a deleterious mutation, the genotypes of both asthenospermic bulls were screened for the presence of long ROH (**Additional files 1 & 2**). We identified 49 and 28 ROH in the two bulls that covered 258 and 210 Mb of their genomes (**Figure 2**). The average length of the ROH was 6.1 Mb ranging from 1.4 to 30.4 Mb. ROH at chromosomes 2, 4, 9, 10, 14 and 25 were present in both asthenospermic bulls. However, the ROH at chromosomes 4, 9, 10 and 14 were homozygous for different haplotypes and thus not compatible with the presumed inheritance of a recessive allele from a common ancestor. Two haplotypes on chromosomes 2 (between 90,155,450 and 92,350,004 bp) and 25 (between 25,490,468 and 28,470,779 bp) were identical-by-descent (IBD) in both bulls with asthenospermia. The IBD haplotype located on chromosome 2 consisted of 28 adjacent SNPs of the BovineSNP50 Bead chip. In 100 randomly chosen fertile bulls of the Nordic Red cattle population, the haplotype had a frequency of 28%. Moreover, the haplotype occurred in the homozygous state in four bulls with normal fertility. Considering its high frequency and occurrence in the homozygous state in fertile bulls, it is unlikely that this haplotype was causal for the asthenospermia of two bulls. The IBD haplotype located on chromosome 25 consisted of 45 adjacent SNPs of the BovineSNP50 Bead chip (**Additional file 6**). It was not detected in the homozygous state in normospermic bulls. Both the sires and ten (out of 17) half-sibs of the asthenospermic bulls carried the haplotype in the heterozygous state. The average calving rate was 0.42 and 0.41 in half-sibs that were haplotype carriers and non-carriers, respectively, indicating that fertility is normal in heterozygous bulls. In 8,557 fertile sires from the Nordic Red cattle genetic evaluation reference population, the haplotype was observed in the heterozygous state in 426 animals corresponding to a frequency of 2.5%. Homozygous haplotype carriers were absent among the fertile bulls although five were expected assuming Hardy-Weinberg Equilibrium proportions (P=0.018), corroborating recessive inheritance of a deleterious allele.

**Figure 2.**
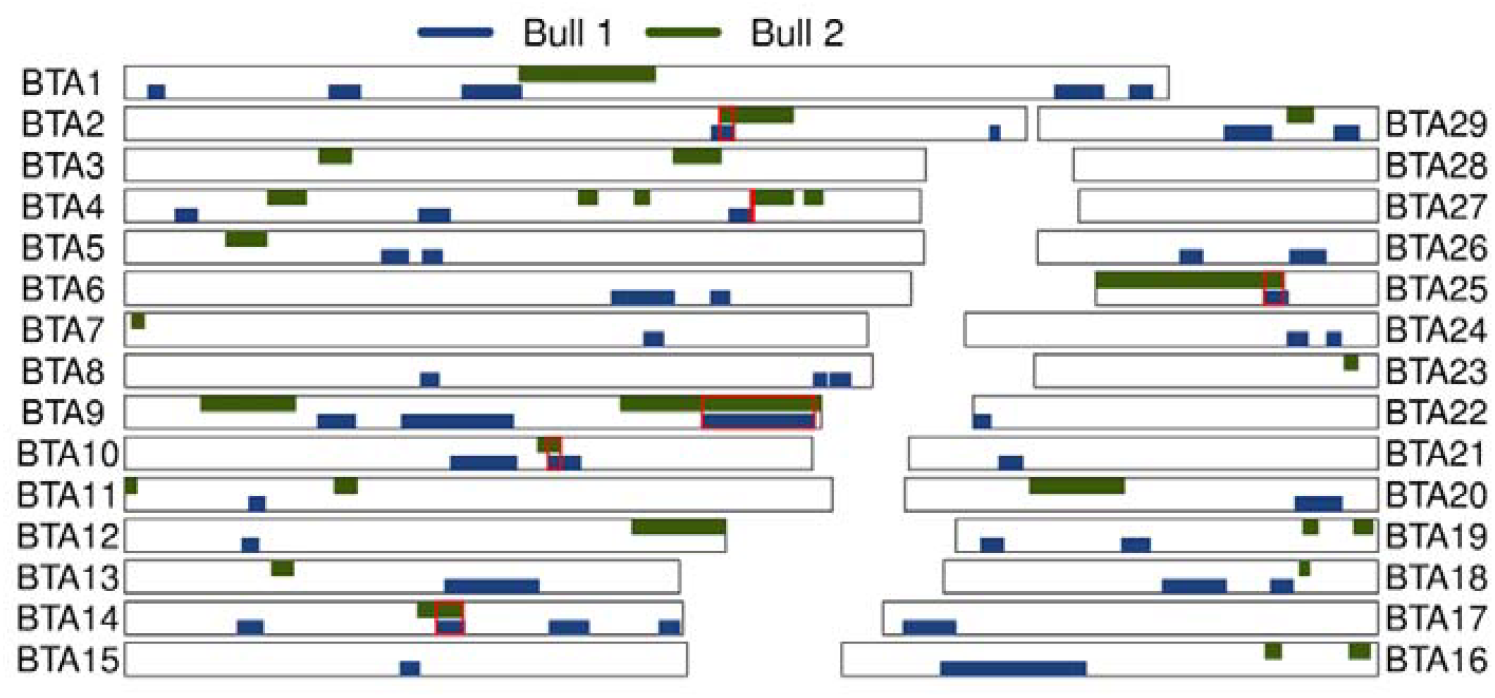
Homozygosity mapping in two asthenospermic bulls. Homozygosity mapping in two asthenospermic bulls. Blue and green color represent autosomal runs of homozygosity (ROH) that have been detected in the two asthenospermic bulls. The red frames highlight six ROH that had been detected in both affected bulls. Only the 2.98-Mb segment located on chromosome 25 was never found in the homozygous state in normospermic bulls.

### A variant disrupting a splice donor site in *CCDC189* is associated with asthenospermia

In order to detect the causal mutation for the asthenospermia, we sequenced both affected bulls at approximately 12-fold genome coverage. The sequencing depth in the interval between 25,490,468 and 28,470,779 bp on chromosome 25, where both bulls shared an extended segment of autozygosity, did not differ from the average sequencing coverage (**Additional file 7**). Visual inspection of the aligned reads in the 2.98 Mb interval revealed no evidence for a large deletion or structural variant that might segregate with the haplotype. We identified 3,683 single nucleotide and short insertion and deletion polymorphisms that were homozygous for the non-reference variant in both bulls in the 2.98 Mb IBD interval. Out of 3,683 sequence variants, 3,657 occurred in the homozygous state in at least one out of 237 male control animals from breeds other than Nordic Red that were not affected by the disorder. Such variants are not compatible with the presumed recessive inheritance of a fully penetrant allele that causes asthenospermia, thus retaining 126 candidate causal variants. Variants annotation using the *Variant Effect Predictor* tool revealed that 120 of 126 candidate causal variants were located in non-coding regions of the genome (**Additional file 8**). Six variants were predicted to affect protein-coding regions or splice sites including four synonymous mutations, a missense variant (GCA_000003055.3:Chr25:g.26184191C>G) in *RABEP2* (ENSBTAT00000008595:p.P413R, SIFT-score [25] 0.26) and a variant disrupting a canonical 5’ splice donor site (GCA_000003055.3:Chr25:g.27138357C>T) in *CCDC189* (transcript-ID: ENSBTAT00000045037.3:c.472+1G>A) encoding the coiled-coil domain containing protein 189. Of the 126 candidate causal variants, only the splice donor site mutation in *CCDC189* was predicted to be deleterious to protein function.

The polymorphism Chr25:27138357C>T alters the first position of the fourth intron of bovine *CCDC189*. Please note that our results are based on the UMD3.1 assembly of the bovine genome. According to the latest assembly and annotation of the bovine genome (ARS-UCD1.2), the polymorphism is located at GCF_002263795.1:Chr25:g.26880841C>T and ENSBTAT00000045037.4:c.472+1G>A. The derived T-allele disrupts a canonical splice site (GT>AT; *CCDC189* is transcribed from the reverse strand, **Figure 3**). The T-allele did not segregate in 237 healthy animals from cattle breeds other than Nordic Red that were sequenced in Schwarzenbacher *et al.* [26]. In 2669 animals that had been sequenced in the framework of the 1000 Bull Genomes Project, the Chr25:27138357 T allele occurred in the heterozygous state only in four animals of the Finnish Ayrshire and Swedish Red cattle breeds (**Additional file 8**), i.e., two breeds that contribute genes to the Nordic Red dairy cattle breed [27]. The T allele was never observed in the homozygous state in cattle other than the two bulls with asthenospermia.

**Figure 3.**
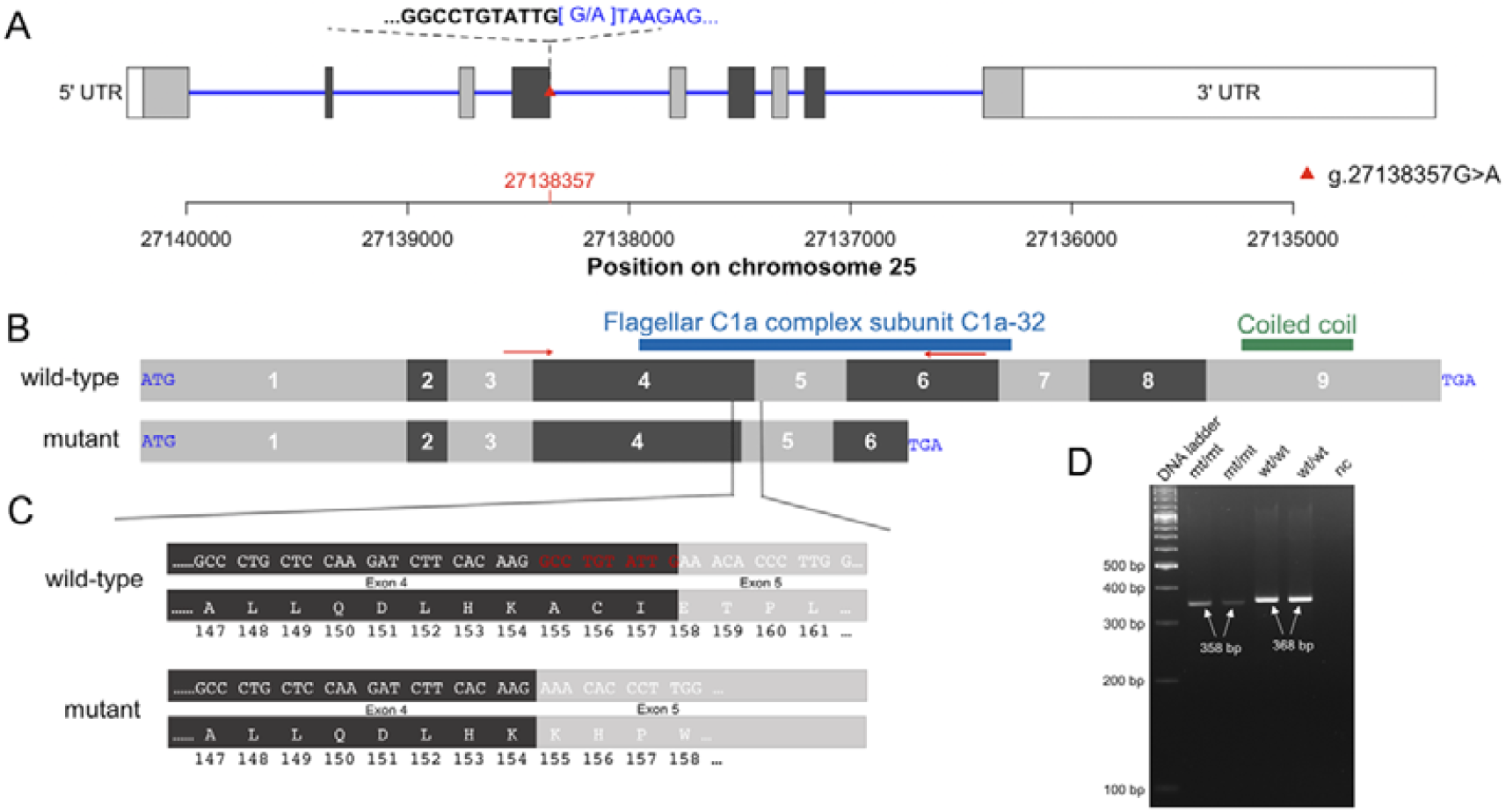
Schematic overview of bovine *CCDC189.* Structure of the bovine *CCDC189* gene (transcript-ID: ENSBTAT00000045037). The bovine *CCDC189* gene encoding the coiled-coil domain containing protein 189 is transcribed from the reverse strand. It consists of nine exons (grey and dark-grey boxes) (A). A mutant allele (red triangle) at 27,138,357 basepairs disrupts a canonical 5’ splice donor variant in the fourth intron. Schematic view of the cDNA sequence of *CCDC189* (B). The coiled-coil domain containing protein 189 contains the flagellar C1a complex subunit C1a-32 between amino acids 127 and 221 and a coiled coil domain between amino acids 280 and 308. Red arrows indicate the positions of the primers used for RT-PCR. The mutant allele activates a cryptic splice site in exon 4 that truncates 10 basepairs from exon 4 (red colors) and introduces a frameshift in translation that eventually creates a premature stop codon (C). The mutant protein, if retained, lacks a large fraction of the flagellar C1a complex subunit C1a-32 and the coiled coil domain. Effect of the mutant allele on CCDC189 transcription (D). The RT-PCR products from testis tissue were shorter from two mt/mt (358 bp) than wt/wt (368 bp) bulls. Lane 5 (nc) is the negative PCR control. The length of the bands was defined using the Image Lab version 5.0 software (Bio-Rad Laboratories).

The effect of the Chr25:27138357C>T variant on *CCDC189* transcription was examined by RT-PCR using RNA extracted from testicular tissues of the two asthenospermic and two normospermic control bulls that were homozygous for the wild type allele. Using primers located at the junction of exons 3 and 4 (forward) and in exon 6 (reverse) we obtained 358 and 368 bp RT-PCR products in asthenospermic (mt/mt) and normospermic (wt/wt) bulls, respectively (**Figure 3D**). Sequence analysis of the RT-PCR product of mt/mt bulls revealed that the Chr25:27138357C>T substitution activates a cryptic splice site in exon 4. It alters the resulting CCDC189 protein sequence from amino acid residue 155 onwards and creates a premature stop codon at amino acid residue 200, thus truncating 40% of the 332-amino acid CCDC189 protein. The truncated protein lacks domains that are supposed to be essential for physiological protein function including the flagellar C1a complex subunit C1a-32, which modulates ciliary and flagellar beating [28].

### Prevalence of the splice donor variant in Nordic Red cattle

In order to confirm the association between the candidate causal splice site mutation in *CCDC189* and the asthenospermia-associated haplotype, we used Sanger sequencing to obtain genotypes for the polymorphism in 118 normospermic control bulls of the Nordic Red cattle breed. The variant was in perfect linkage disequilibrium with the 2.98 Mb haplotype corroborating that it might be causative for the disorder. All bulls carrying the associated haplotype in the heterozygous state were also heterozygous for the Chr25:27138357 T allele. The T allele was not found in bulls that did not carry the asthenospermia-associated haplotype.

## Discussion

A splice donor variant in bovine *CCDC189* is the likely cause of asthenospermia in the Nordic Red cattle breed. An autosomal recessive mode of inheritance was suspected because the two affected bulls were related through a common ancestor on both their paternal and maternal paths, the spermatological findings were very similar in their ejaculates, and their first-and second-degree relatives were fertile. Increasing rates of inbreeding and the widespread use of individual sires in artificial insemination make cattle populations susceptible to the propagation and phenotypic manifestation of recessive alleles [29],[30]. Considering that only two bulls of the breed were affected by asthenospermia, explanations other than recessive inheritance were also plausible including, e.g., allelic heterogeneity [31],[32] and non-genetic causes of disease. However, both bulls were housed at the AI center under controlled environmental conditions at the same time as many fertile bulls and their health and andrological parameters were normal indicating that non-genetic factors were less likely to cause this condition. Previous studies showed that autozygosity mapping is a powerful approach to identify genomic regions associated with recessive conditions even if only few affected individuals with genotype information are available [13],[33]. Using autozygosity mapping in the two affected bulls, we identified a 2.98 Mb segment of extended homozygosity located at chromosome 25 that was identical by descent in both bulls. This segment, encompassing a variant disrupting a canonical 5’ splice donor site in *CCDC189*, was never found in the homozygous state in fertile bulls, in agreement with the presumed recessive inheritance of an allele with detrimental phenotypic consequences. Two additional bulls from the Nordic Red cattle breed were reported to us in retrospect from technicians of an AI center, because they produced ejaculates with immotile sperm. Both bulls were genotyped for genomic evaluation using bovine 50K SNP arrays. It turned out that these two bulls were also homozygous for the 2.98 Mb segment on chromosome 25 further corroborating that this segment harbors a recessively inherited causal mutation for asthenospermia (**Additional file 9**). Unfortunately, neither DNA, nor semen nor or any other tissue of those bulls has been preserved for functional investigations.

Verifying the effect of candidate causal mutations in independent populations is critical evidence to supporting variant causality [34],[26]. The splice donor variant in *CCDC189* did not segregate in breeds distantly related to Nordic Red cattle indicating that it likely arose after the formation of breeds thus precluding an immediate validation of the mutation in an unrelated cattle population. Our study thus exemplifies difficulties inherent in validation and functional investigation of deleterious sequence variants that manifest severe phenotypic consequences such as impaired male fertility in livestock populations. Due to purifying selection, deleterious sequence variants are rare in natural populations [34] unless they either show pleiotropic effects on desired traits [31], are in linkage disequilibrium with quantitative trait loci [35], or are propagated through the widespread use of heterozygous mutation carriers in AI [36],[4]. The splice donor variant in *CCDC189* is rare in the Nordic Red dairy cattle breed. We detected the derived allele in its homozygous state in only two bulls for which tissue was available for functional investigations. Assuming an allele frequency of the derived allele of 2.5% and random mating, only 1 out of 1,600 males is expected to be homozygous and thus sterile due to spermatozoa with impaired motility. The disorder remains undetected in most homozygous animals because sterility becomes evident only in breeding bulls. Apart from producing asthenospermic ejaculates, both bulls homozygous for the derived T allele were healthy. However, genetic variants that cause asthenospermia may also compromise the function of motile cilia thus manifesting themselves in chronic detrimental diseases [11],[37],[26]. Further research is warranted to determine if DNA variants in *CCDC189* are associated with ciliopathies particularly in animals that are kept in conventional farms under normal husbandry conditions. A direct gene test will facilitate detection of and selection against carriers in order to prevent the birth of homozygous males that are sterile. Due to the low frequency of the deleterious allele, negative effects on genetic gain and genetic diversity are likely negligible when carrier animals are excluded from breeding in the Nordic Red cattle breed. However, sophisticated genome-based mating strategies are required to take into account the steadily increasing number of deleterious sequence variants discovered in breeding populations [38].

Functional and pathophysiological investigations were possible in two AI bulls only. Sperm analysis revealed an identical pattern in both bulls homozygous for the deleterious allele; sperm motility was impaired in both fresh ejaculates and post-thaw, e.g., the process of handling the semen, even with dilution of the seminal plasma did not ameliorate the low progressive motility. Slightly elevated ATP concentrations in post-thaw semen of the affected bulls might result from lower sperm metabolism or accumulation of ATP due to impaired progressive motility. However, elevated ATP concentrations might also derive from the inactive sperm tails. Moreover, a normal proportion of viable albeit asthenospermic spermatozoa resulted in high superoxide and reactive oxygen species production indicating defective mitochondria [39]. This further impairs sperm motility and eventually leads to lipid peroxidation and DNA damage and/or cell death, thus precluding fertilization [40]. Our study showed that the affected bulls were infertile *in vivo* and *in vitro*, precluding their use in AI. Fertilization might be possible *in vitro* using intracytoplasmic sperm injection, though DNA damage could impair embryo development. We have not explored this possibility, since the sperm phenotypic deviations were decisive to cull breeding sires from AI.

Most likely the causal variant for asthenospermia disrupts a canonical 5’ splice donor site of bovine *CCDC189* encoding the coiled-coil domain containing protein 189. To the best of our knowledge, this is the first time that a phenotypic consequence has been associated with a variant of *CCDC189* in any organism. CCDC189 (alias Spergen-4) is a sperm tail-associated protein that is mainly located in the principal piece and acrosome region of mature spermatozoa indicating that it may be required for their physiological activated progressive movement and acrosome reaction [41]. The velocity of spermatozoa was slower and their progressive motility was severely compromised in bulls that were homozygous for the Chr25:27138357 T allele indicating that flagellar movement was defective. Mutations disrupting components of the dynein arms or radial spokes of axonemal dyneins cause disorganized axonemes of motile cilia and flagella, thus compromising their physiological movement [11],[42],[43]. In spite of an impaired motility, the ultrastructural organization of the sperm flagella including the assembly of the dynein arms was normal in both asthenospermic bulls, indicating that disorganized axonemes were less likely to be responsible for the lack of progressive movement of the sperm. We showed that the Chr25:27138357 T allele introduces a cryptic splice site in the fourth exon of *CCDC189* leading to a frameshift in translation and premature translation termination. Considering that the mutant transcript including the premature stop codon is likely degraded via nonsense-mediated mRNA decay or, if not degraded via nonsense-mediated mRNA decay, would lack a large part of the flagellar C1a complex subunit (C1a-32), that is an integral constituent of the central pair of flagellar microtubules, the splice donor site of *CCDC189* most likely represents a loss-of-function variant. Aberrations of the small C1a microtubule projection of the central pair impair the physiological beating of flagella whose axonemes are normally organized otherwise [44],[45]. It has been shown that the C1a-32 subunit mediates the beating of cilia and flagella by modulating the activity of both the inner and outer dynein arms [28],[46]. Possibly, the resolution of the transmission electron microscopy was too low to reveal potential aberrations of the small C1a-32 subunit of the C1a projection [28]. Both bulls homozygous for the splice donor variant were sterile because they produced asthenospermic ejaculates corroborating that CCDC189 is important for physiological movement of spermatozoa and fertilization. However, spermatozoa of both asthenospermic bulls bended slightly which is not classified as progressive motility in automated semen analysis. The minor movement of affected spermatozoa resembles uncoordinated twitching of flagella in *Chlamydomonas* mutants lacking constituents of the flagellar C1a projection [45].

## Conclusions

Our study in cattle revealed that a naturally occurring deleterious allele of *CCDC189* encoding the coiled-coil domain containing protein 189 causes immotile sperm with defective mitochondria thus resulting in male sterility both *in vivo* and *in vitro*. The mutation occurs in Nordic Red dairy cattle at a low frequency. Direct gene testing facilitates the efficient monitoring of the deleterious allele in the population.

## Material and Methods

### Animals

Two fifteen-months old bulls from the Nordic Red dairy cattle breed that were born in 2012 and 2013 were housed at an AI center in order to collect their semen for AI. Both bulls were apparently healthy and their andrological status was considered as normal, i.e., they showed no signs of disease, the size of their testicles was within the normal range and their libido was normal. During a period of more than four months, 11 and 27 ejaculates, respectively, of both bulls were collected once or twice a week by technicians at the AI center. Semen from bulls of documented normal sperm quality and fertility, of the same breed and age interval, was likewise collected, processed and examined. Testicular tissue samples were collected immediately after slaughter, frozen in liquid nitrogen and stored at −80°C.

### Analysis of fresh semen

Immediately after semen collection, sperm motility, viability and morphology of fresh ejaculates were subjectively assessed by technicians at the AI center. Sperm motility was evaluated using a microscope equipped with phase contrast optics. Sperm concentration and plasma membrane viability were assessed using a flow-cytometer [47]. None of the collected ejaculates fulfilled the requirements for AI because most spermatozoa had reduced or missing progressive motility. Upon our request, one ejaculate from each of the asthenospermic bulls was frozen using the commercial method routinely used by the AI center.

### Analysis of frozen-thawed semen samples

Velocity (µm/sec), and percentages of spermatozoa with total and progressive motility were assessed using frozen-thawed ejaculates. Semen droplets (2.4 × 10^5^ spermatozoa in 10 µL) were placed on a pre-warmed 76 × 26 mm Menzel-Gläser microscope slide (ThermoFisher Scientific, Waltham, MA, USA) covered by a pre-warmed coverslip on a thermal plate (38 °C). Slides were analyzed using an upright Zeiss Axio Scope A1 light microscope equipped with a 10x phase contrast objective (Carl Zeiss, Stockholm, Sweden) connected via a CMOS camera (UEye, IDS Imaging Development Systems GmbH, Obersulm, Germany) to a computer running the Qualisperm™ software (Biophos SA, Lausanne, Switzerland). The Qualisperm™ technology is based on fluorescence correlation spectroscopy-analysis of single particles (spermatozoa) in confocal volume elements. Individual spermatozoa are projected on a pixel grid of the CMOS camera and the algorithm calculates the number of fluctuations in each pixel using a correlation function of sperm numbers and translation classes, thus determining the speed (velocity) distribution of spermatozoa, from which proportions of motility classes are calculated [48].

Sperm viability (membrane integrity) and early membrane destabilization changes (YO-PRO-1+/PI-), degree of mitochondrial activity (Mitotracker Deep Red+) and superoxide production (MitoSOX+), production of reactive oxygen species (ROS; CellROX+) and lipid peroxidation (BODIPY+) were assessed using a Gallios™ flow cytometer (Beckman Coulter, Bromma, Sweden) equipped with standards optics. In all investigations, we assessed 25,000 events per sample with a flow rate of 500 cells per second.

The total ATP concentration of frozen-thawed spermatozoa was analyzed using the ATP Determination Kit A22066 (Molecular Probes TM Invitrogen detection technologies) according to the manufacturer’s protocol. Samples of an asthenospermic and normospermic bull were analyzed using a Kikkoman Lumitester C-100 luminometer (Kikkoman Corporation, Nishishinbashi, Tokyo, Japan).

Transmission electron microscopy on spermatozoa of two asthenospermic and one normospermic bull was performed using frozen-thawed semen samples. After thawing, semen samples were washed twice in phosphate buffered saline (PBS), fixed with 2.5% glutaraldehyde and 2% paraformaldehyde in PBS for 2 hours on ice and pelleted at 6000g. Pellets were washed in PBS, post-fixed in 1% osmium tetroxide, washed in ddH_2_O, stained in 1% uranyl acetate, dehydrated in graded series of ethanol (25%, 50%, 75%, 90%, and 100%), and embedded in Epoxy resin through increasing concentrations (25%, 50%, 75%, and 100%) using PELCO Biowave+ tissue processor, and then cured at 60°C for 3 days. Embedded blocks were sectioned using Leica FC6 microtome and a DIATOME diamond knife with 45° angle into 60nm sections and mounted on Quantifoil copper grids with formvar carbon films. Sections were post-stained with 2% uranyl acetate followed by lead citrate. Grids were imaged using FEI Morgani 268 electron microscope operated at 100 kV at 20k magnification.

### *In vivo* and *in vitro* fertilization success

Fertility of two asthenospermic bulls was tested *in vivo* at a conventional Finnish cattle herd within one breeding season; ninety-eight cows in oestrus were inseminated by the farmer either with semen from one of two bulls with asthenospermia (n=9) or a control bull with normal semen quality (n=89). Insemination success was controlled by a trained veterinarian.

*In vitro* fertilization (IVF) success, with embryo cleavage rate at 42 hours post insemination and embryo yield at 9 days post insemination as end-points, was tested for an asthenospermic and a normospermic bull, using 470 offal oocytes. Ovaries of slaughtered cows were collected at a commercial slaughterhouse and transported to the laboratory in saline at room temperature. In the laboratory, follicles of 2-8 mm in diameter were aspirated with an 18 G needle and a 20 ml syringe. Cumulus-oocyte complexes (COC) were collected from the aspirated follicular fluid and 470 good quality COC were selected for maturation. Unless stated otherwise, all the chemicals were purchased from Sigma-Aldrich (St. Louis, MO, USA). COC were matured for 24h in TCM199 with glutamax-I (Gibco™; Invitrogen Corporation, Paisley, UK) supplemented with 0.25mM Na-pyruvate, 100 IU/ml penicillin, 100 μg/ml streptomycin, 2 ng/ml FSH (Puregon, Organon, Oss, Netherlands), 1 μg/ml β-estradiol (E-2257) and 10% heat inactivated fetal bovine serum (FBS) (Gibco™, New Zealand) at 38.5°C in maximal humidity in 5% CO_2_ in air. Following maturation, 320 and 150 COC were inseminated using washed frozen–thawed semen from an asthenospermic and a normospermic control bull of proven fertility in IVF, respectively, at a concentration of 1 million spermatozoa per ml at 38.5°C in maximal humidity in 5% CO_2_ in air for 20h. During IVF spermatozoa capacitation and movement were stimulated with 10 μg/ml heparin and PHE (20 µM penicillamine – 10 µM hypotaurine – 1 µM epinephrine), respectively. After fertilization, presumptive zygotes were cultured in G1/G2 media (Vitrolife, Göteborg, Sweden) supplemented with fatty-acid-free bovine serum albumin (4 mg/ml) at 38.5°C in maximal humidity in an atmosphere of 5% O_2_ and 5% CO_2_. Cleavage rates of oocytes were recorded at 30 and 42 hours post insemination. The number and quality of blastocysts were recorded at day 8 and 9 (IVF=day 0).

### Genotyping, quality control and haplotype inference

The two asthenospermic bulls were genotyped at 54,001 SNPs using the Illumina BovineSNP50 Bead chip using DNA purified from semen samples following standard DNA extraction protocols. To facilitate haplotype inference, we also considered genotypes of 119 fertile bulls from the Nordic Red dairy cattle breed, including the sires (n=2) and 17 paternal half-sibs of the affected bulls. Genotypes for these 119 bulls were also determined using the Illumina BovineSNP50 Bead chip comprising genotypes at 54,001 (version 1) or 54,609 SNPs (version 2). The chromosomal positions of the SNPs corresponded to the University of Maryland reference sequence of the bovine genome (UMD3.1) [49]. Following quality control, we retained 41,594 autosomal SNPs with a call-rate greater than 95%. The per-individual call-rate was between 96.07% and 100% with an average call-rate of 99.73%. The *Beagle* [50] software (version 3.2.1) was used to impute sporadically missing genotypes and infer haplotypes. Genotype data are available in *plink* binary format [51] (.bed/.bim/.fam) as **Additional file 1**.

### Detection of runs of homozygosity

Runs of homozygosity (ROH) were identified in both asthenospermic bulls using the *homozyg*-function that was implemented in the *plink* (version 1.9) software tool [51]. Due to the relatively sparse SNP coverage (approximately 1 SNP per 60 kb), we considered segments with a minimum length of 500 kb *(--homozyg-kb 500*) and at least 20 contiguous homozygous SNPs (*--homozyg-snp 20*) as ROH. ROH were identified within sliding windows of 40 SNPs (*--homozyg-window-snp 40*). For the identification of ROH, no heterozygous SNP were allowed within ROH (*--homozyg-het 0*). The R script that was used to identify and visualize ROH is available as **Additional file 2**.

### Identification of segments of autozygosity in two bulls with asthenospermia

Genotypes contained in ROH in both asthenospermic bulls were manually inspected in order to determine whether the bulls were identical by descent (IBD) for the same haplotype (i.e., autozygous). In regions where both affected bulls were IBD, the haplotypes of 119 fertile bulls (see above) were also inspected. Assuming recessive inheritance of a deleterious mutation, we considered IBD haplotypes as associated with asthenospermia if they were not detected in the homozygous state in 119 fertile bulls.

### Generation of whole genome sequencing data and sequence variant detection

Genomic DNA was prepared from semen samples of the two asthenospermic bulls according to standard DNA extraction protocols. Paired-end libraries were prepared using the paired-end TruSeq DNA sample preparation kit (Illumina) and sequenced using a HiSeq 2500 instrument (Illumina). We collected 252 and 270 million paired-end reads (125 base pairs) corresponding to an average (raw) genome coverage of 12- and 13-fold, respectively. The raw sequencing data of both asthenospermic bulls are available at the European Nucleotide Archive (ENA) (http://www.ebi.ac.uk/ena) under primary accession number PRJEB29487. The raw sequencing reads were trimmed and filtered using *fastp* (version 0.19.4) [52] and subsequently aligned to the UMD3.1 reference sequence (GCA_000003055.3) of the bovine genome [49] using the *BWA mem* algorithm (version 0.7.13) [53]. Individual files in SAM format were converted into BAM format using *SAMtools* (version 1.8) [54]. Duplicate reads were marked with the *MarkDuplicates* command of the *Picard tools* software suite (https://broadinstitute.github.io/picard/). Polymorphic sequence variants were identified using the *Genome Analysis Toolkit* (*GATK*, version 4.0) [55]. Sequencing coverage along the genome was calculated based on the aligned reads using the *Mosdepth* software [56].

### Identification of candidate causal variants

Assuming that both bulls with asthenospermia were homozygous for the same recessively inherited variant, we screened the sequence data of the affected bulls for variants that were: a) located within the 2.98-Mb segment of autozygosity, b) homozygous for the non-reference allele in both bulls and c) never observed in the homozygous state in a control group of 237 animals from breeds other than Nordic Red dairy cattle that were obtained from Schwarzenbacher *et al.* [26]. 126 variants that fulfilled these three criteria were considered as candidate causal variants. The functional consequences of the candidate causal variants were predicted according to the Ensembl transcripts database (release 91) using the *Variant Effect Predictor* (VEP) [57] tool. Genotypes of candidate causal variants were subsequently inspected in whole-genome sequencing data of 2,669 animals that have been sequenced in Run6 of the 1000 Bull Genomes Project [58].

### Validating the CCDC189 splice donor variant in Nordic Red cattle

We estimated the frequency of the asthenospermia-associated haplotype in 8,557 Nordic Red cattle from the Nordic genomic selection reference population that had been genotyped at more than 50,000 SNPs [27]. Using Sanger sequencing, we genotyped the splice donor variant (GCA_000003055.3:Chr25:g.27138357C>T) in *CCDC189* (transcript-ID: ENSBTAT00000045037) in 118 sires that were either identified as heterozygous carriers of the asthenospermia-associated haplotype (N=30) or have been used frequently in AI (N=88). PCR reactions for genotyping were done with the DyNAzyme II DNA Polymerase (Thermo Fisher, MA, US) in a 10µl volume of 1x PCR buffer, 0.2mM dNTPs, 10pmol primer mix (forward 5’-CAAGGTCCTGCCCTTAAGAA-3’) and reverse primer: (reverse 5’-ATGCCATCACTCTGGACCTC-3’) and 150pg of DNA. The cycling conditions were the following: a) an initial denaturation at 95°C for 3 min, b) 29 cycles of 30 sec denaturation (94°C), 30 sec hybridization (58°C), 30 sec elongation (72°C), and c) a final 3 min elongation (72°C). PCR products were purified and directly sequenced using the BigDye Terminator Cycle Sequencing Kit (Applied Biosystems, CA, US). Electrophoresis of sequencing reactions was performed on 3500xL Genetic Analyzers (Applied Biosystems, CA, US), and sequences were visualized with Sequencher 5.4.6 (Gene Codes Corporation, MI, US).

### Transcription analysis

Total RNA from two bulls homozygous for the *CCDC189* splice donor variant (GCA_000003055.3:Chr25:g.27138357C>T) and two control bulls that were homozygous for the wild type allele was extracted from testis tissues using the RNeasy Mini Kit (Qiagen) following manufacturer’s instructions. The quality and concentration of total RNA was analysed using Agilent’s 2100 Bioanalyzer and the Nanodrop ND-2000 spectrophotometer (Thermo Scientific). An amount of 500 ng of total RNA from each sample was reversely transcribed using QuantiTech Rev. Transcription kit (Qiagen). PCR reactions were done with the DyNAzyme II DNA Polymerase (Thermo Fisher, MA, US) in a 30 µl volume of 1 × PCR buffer, 0.2 mM dNTPs, 10 pmol primer mix (forward 5’-ACTGAGGAGATGAGGGAGGT-3’ and reverse primer: reverse 5’-TAGAGCTTGAAGTGGCGGAA-3’) and 50 ng of cDNA. The cycling conditions were the following: 1) an initial denaturation at 95°C for 3 min, 2) 35 cycles of 30 sec denaturation (94°C), 30 sec hybridization (61.8°C), 1 min elongation (72°C) and 3) a final 3 min elongation (72°C). PCR products were separated on a 4% agarose gel and the length of the products was analysed using the Image Lab version 5.0 software (Bio-Rad Laboratories). PCR products were purified with QIAquick Gel Extraction kit (QIAgen) and sequenced using the BigDye Terminator Cycle Sequencing Kit (Applied Biosystems, CA, US). Electrophoresis of sequencing reactions was performed on 3500xL Genetic Analyzer (Applied Biosystems, CA, US), and sequences were visualized with Sequencher 5.4.6 (Gene Codes Corporation, MI, US).

## Declarations

### Ethics approval and consent to participate

Ejaculates of two asthenospermic bulls were collected by employees of an AI center in Finland as part of their routine practice under veterinary supervision. Both bulls were eventually culled because their impaired sperm motility precluded the use of the collected ejaculates for artificial inseminations. The decision to cull the bulls was at the sole discretion of representatives of the AI center. Testicular tissue was collected immediately after slaughter. Oocytes used for *in vitro* fertilization experiments were collected from offal cow ovaries at a commercial slaughterhouse. No animal experiments were performed in this study, and, therefore, approval from an ethics committee was not required.

### Consent for publication

Not applicable

### Availability of data and material

Whole-genome sequence data of both asthenospermic bulls have been deposited at the European Nucleotide Archive (ENA) of EMBL-EBI under sample accession numbers SAMEA5064546 and SAMEA5064547 at primary accession PRJEB29487 (https://www.ebi.ac.uk/ena/data/view/PRJEB29487). Genotype data of two asthenospermic bulls are available in *plink* binary format as supporting information. Whole-genome sequence data of all other animals used in this study are part of the 1000 Bull Genome Project. Parts of the whole-genome sequence data of individual bulls of the 1000 Bull Genomes Project (http://www.1000bullgenomes.com/) are available at https://doi.org/10.1038/s41588-018-0056-5 and the European Nucleotide Archive (ENA) under the primary accession numbers PRJNA238491, PRJEB9343, PRJNA176557, PRJEB18113, PRJNA324822, PRJNA324270, PRJNA277147, PRJEB5462. The BovineSNP50 Bead chip genotypes of all other animals included in this study have been collected for routine genomic evaluations and are part of the Nordic Cattle Genetic Evaluation genotype database (https://www.nordicebv.info).

### Authors’ contributions

Sample collection: HV, MA; Analysis of genotyping data: TIT HV AS GS BG HP; Analysis of whole-genome sequencing data: CW RF HP; RNA analysis: TIT AS; Sperm functional analyses: AVC MAR SN HS HRM MA MH HV KFi KFl TS; In vitro fertilization experiment: JP MM; In vivo fertilization experiment: JT; Transmission electron microscopy: MH; Wrote the paper: TIT MA HRM HP; Conceived and designed the experiments: HV MA HP; Read and approved the final version of the manuscript: all authors

## Acknowledgements

Genotype data of the control bulls were kindly provided by Viking Genetics. The authors thank the Scientific Center for Optical and Electron Microscopy (ScopeM) of ETH Zurich, particularly Dr. Cecilia Bebeacua for support in transmission electron microscopy.

## Supplementary Information

### Additional file 1 File S1

File format: .zip

Title: Genotypes of two asthenospermic bulls

Description: Genotypes at 41,594 SNPs for two asthenospermic bulls in plink binary format. The positions of the SNP correspond to the UMD3.1 assembly of the bovine genome.

### Additional file 2 File S2

Format: .txt

Title: R script to identify IBD segments in two asthenospermic bulls

Description: R script to visualize runs of homozygosity in the genotype data of two asthenospermic bulls. This program uses as input the files provided as Supplementary File 1. Please note that this R script calls the fast algorithm implemented in plink (version 1.9) to identify runs of homozygosity.

### Additional file File S3

Format: .mov

Title: Post-thaw sperm motility in an asthenospermic bull

Description: Video of frozen-thawed spermatozoa of an asthenospermic bull. Please note the minor movement of some spermatozoa that is typically not classified as progressive motility during the subjective eye-evaluation of ejaculates.

### Additional file 4 File S4

Format: .mov

Title: Post-thaw sperm motility in a normospermic control bull

Description: Video of frozen-thawed spermatozoa of an normospermic bull.

### Additional file 5 File S5

Format: .png

Title: Pedigree of two asthenospermic bulls

Description: Pedigree of two affected bulls. Ovals and boxes represent female and male ancestors, respectively. The grey box highlights the most recent common ancestor (MRCA) that was present on both the maternal and paternal ancestry of both asthenospermic bulls.

### Additional file 6 File S6

Format: .csv

Title: Asthenospermia-associated haplotype

Description: ID, position and asthenospermia-associated allele of 45 adjacent SNPs of the BovineSNP50 Bead chip. The coding of the alleles is based on the Illumina TOP/BOT format. The chromosomal position of the SNP corresponds to the UMD3.1 assembly of the bovine genome.

### Additional file 7 File S7

Format: .png

Title: Sequencing depth along chromosome 25 in two asthenospermic bulls Description: The depth of sequencing coverage in two asthenospermic bulls was calculated at bovine chromosome 25 in sliding windows of 5,000 base pairs using the aligned reads. Grey background indicates the segment of extended homozygosity. The red line represents the position of the candidate causal variant-

### Additional file 8 File S8

Format: .txt

Title: Candidate causal variants compatible with recessive inheritance of asthenospermia Description: Functional consequences of 126 candidate causal variants that were compatible with recessive inheritance of asthenospermia, i.e., they were homozygous for the alternate allele in two asthenospermic bulls and either heterozygous or homozygous for the reference allele in 237 healthy control animals. Variants were annotated using the *Variant Effect Predictor* tool. The quality of the sequence variant genotypes for both asthenospermic bulls is shown in columns 7 and 8. The number of heterozygous control animals (out of 237) is shown in column 9. The last column indicates the genotype distribution of 126 candidate causal variants (homozygous for the reference allele | heterozygous | homozygous for the alternate allele) in 2,669 animals of the 1000 bull genomes dataset.

### Additional file 9 File S9

Format: .png

Title: Homozygosity mapping in four asthenospermic bulls

Description: Homozygosity mapping in four asthenospermic bulls (A). Bulls 1 and 2 were already reported in the main part of the manuscript. Bulls 3 and 4 were additionally reported to us because they also produced immotile sperm. However, tissue or semen was not available for bulls 3 and 4. Blue and green color represent runs of homozygosity (ROH) that have been detected in four asthenospermic bulls. The red frame highlights the 2.98-Mb segment located on chromosome 25 that was never found in the homozygous state in normospermic bulls and that carries the splice donor variant in *CCDC189*. Autozygosity mapping on chromosome 25 (B). Blue and pale blue represent homozygous genotypes (AA and BB), heterozygous genotypes (AB) are displayed in light grey. White color indicates missing genotypes. The red bar indicates a common 2.98-Mb segment that was identical by descent in all four bulls.

