## Supporting_Files for "A splice donor variant in *CCDC189* is associated with asthenospermia in Nordic Red dairy cattle": File_S2.R

```
#+++++++++++++++++++++++++++++++++++++++++++++++++++++++++++++++++++++
# R script to identify and visualize IBD segments in dense genotype data
#
# 01.11.2018, Hubert Pausch, ETH Zürich
#
#+++++++++++++++++++++++++++++++++++++++++++++++++++++++++++++++++++++


#+++++++++++++++++++++++
plink_executable <- "/usr/local/bin/plink_v1.9"	# path to the plink (version 1.9) executable
plink_binary_files <- "astheno_case"	# path to binary genotype files for two asthenopermic bulls
#+++++++++++++++++++++++

#+++++++++++++++++++++++
#	create the chromsome skeleton for the plot
#+++++++++++++++++++++++
bim <- read.table(paste(plink_binary_files, ".bim", sep=''))
chrs <- c(1:29); max_length <- numeric()
for (chr in chrs){
	sub <- bim[bim[,1]==chr,]
	max_length <- c(max_length, max(sub[,4]))
	}

	
#xlimit <- ceiling(sum(max_length[c(2, 29)])/1000000)+1
xlimit <- 190
mat <- matrix(0, 15, 5)
colnames(mat) <- c("plot_position", "chr_set1", "chr_set2", "end_set1", "start_set2")
mat[,1] <- 15:1
mat[,2] <- 1:15
mat[,3] <- 30:16
mat[,4] <- max_length[1:15]/1000000
mat[,5] <- xlimit - max_length[30:16]/1000000

#postscript(paste('Homozygosity_mapping.eps'), paper="special",width=10.5,height=6, bg = 'white', horizontal=TRUE)

#+++++++++++++++++++++++
#	plot the chromosome skeleton
#+++++++++++++++++++++++

par(mar=c(5,5,4,5))
plot(1, xlim=c(0, xlimit), ylim=c(1,16), type='n', axes=FALSE, xlab='', ylab='')
rect(c(0), mat[,1]+.1, mat[,4], mat[,1]+.9, col="white", border="grey20")
rect(c(xlimit), mat[2:15,1]+.1, mat[2:15,5], mat[2:15,1]+.9, col="white", border="grey20")
axis(2, at=c(1.5:15.5), labels = paste("BTA", c(15:1), sep=''), las=1,hadj=.25, tick=FALSE, cex.axis=1.3)
axis(4, at=c(1.5:14.5), labels = paste("BTA", c(16:29), sep=''), las=1,hadj=0.7, tick=FALSE, cex.axis=1.3)
par(xpd=TRUE);legend(40,17.8, lwd=6, col=c('#1F407A', '#3C5A0F'), c("Bull 1", "Bull 2"), bty='n', pt.cex=1.4,cex=1.4, ncol=3);par(xpd=FALSE)


#+++++++++++++++++++++++
anims = c("bull_1", "bull_2")	# IDs of the animals as in the plink.fam file
pos_anims <- c(0, 0.43)	# Position of the IBD "lines"
col_anims <- c("#1F407A", "#3C5A0F")
#++++++++++++++++++++++++


#+++++++++++++++++++++++
#	loop through all autosomes and plot IBD segments
#+++++++++++++++++++++++
counts <- c(0, 0)
genome_homozygosity <- c(0, 0)
all_hom_segs <- c()
segs_affected <- 0
segs <- 0
chrs <- c(1:29)
for (chr in chrs)
	{
	system(paste(plink_executable, " --silent --cow --bfile ", plink_binary_files, " --chr ", chr, " --homozyg-snp 20 --homozyg-density 300 --homozyg-kb 500 --homozyg-window-snp 40 --homozyg --homozyg-het 0", sep=""))
	aa <- read.table("plink.hom", header=TRUE)
	segs <- segs+nrow(aa)
	segs_affected <- segs_affected + nrow(aa[is.element(aa[,2], anims),])
	for (j in 1:length(anims))
		{
		bb <- aa[is.element(aa[,2], anims[j]),]
		if (nrow(bb) >= 1)
			{
			counts[j] <- counts[j] + nrow(bb)
			genome_homozygosity[j] <- genome_homozygosity[j] + sum(bb$POS2-bb$POS1)
			all_hom_segs <- c(all_hom_segs, bb$POS2-bb$POS1)
			if (chr <= 15)
				{
				plot_position_y <- mat[mat[,2]==chr, 1]
				plot_position_x <- 0
				}
			if (chr > 15)
				{
				plot_position_y <- mat[mat[,3]==chr, 1]
				plot_position_x <- mat[mat[,3]==chr, 5]
				}
			rect(plot_position_x+(bb$POS1/1000000), 0.1 + plot_position_y + pos_anims[j], plot_position_x+bb$POS2/1000000,  0.1 + plot_position_y + pos_anims[j]+0.333, col=col_anims[j], border = col_anims[j], lend=1)
			}
		}
	cat(paste("BTA ", chr, " done\n", sep=""))
	}


#+++++++++++++++++++++++
#	highlight regions that are IBD in both bulls
#+++++++++++++++++++++++
rect(90.155450, 14.1, 92.350004, 15-.1, col="NA", border="red2", lwd=1.5) # chr2
rect(94.902406, 12.1, 95.336471, 13-.1, col="NA", border="red2", lwd=1.5) # chr4
rect(87.497282, 7.1, 104.560004, 8-.1, col="NA", border="red2", lwd=1.5) # chr9
rect(64.159733, 6.1, 66.090075, 7-.1, col="NA", border="red2", lwd=1.5) # chr10
rect(47.194767, 2.1, 51.285167, 3-.1, col="NA", border="red2", lwd=1.5) # chr14
rect(147.2008+25.490468, 10.1, 147.2008+28.470779, 11-.1, col="NA", border="red2", lwd=1.5) # chr25

#dev.off()
```
