## Supplementary figures and images for "A splice donor variant in *CCDC189* is associated with asthenospermia in Nordic Red dairy cattle"

### File_S5.png

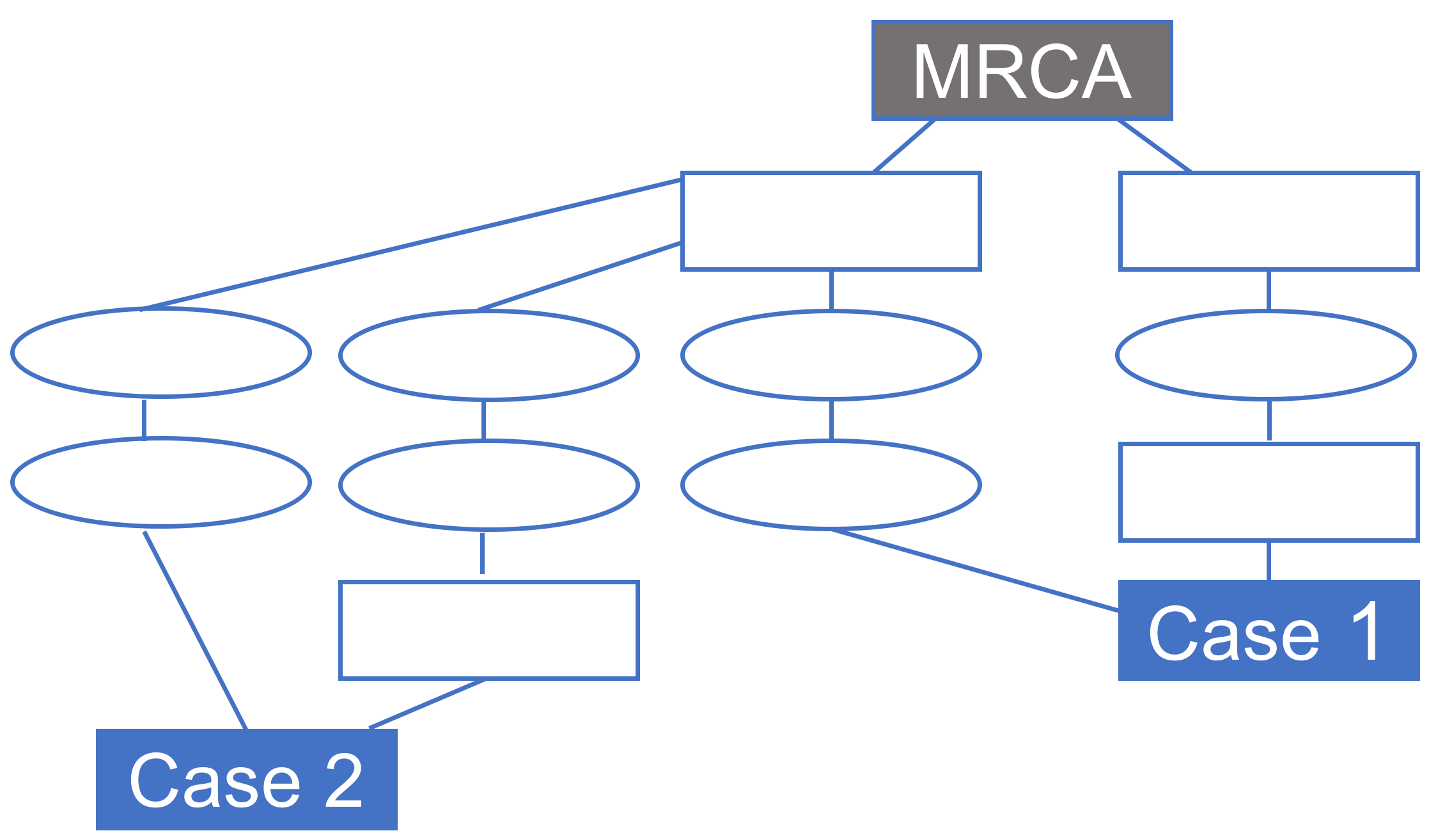

### File_S7.png

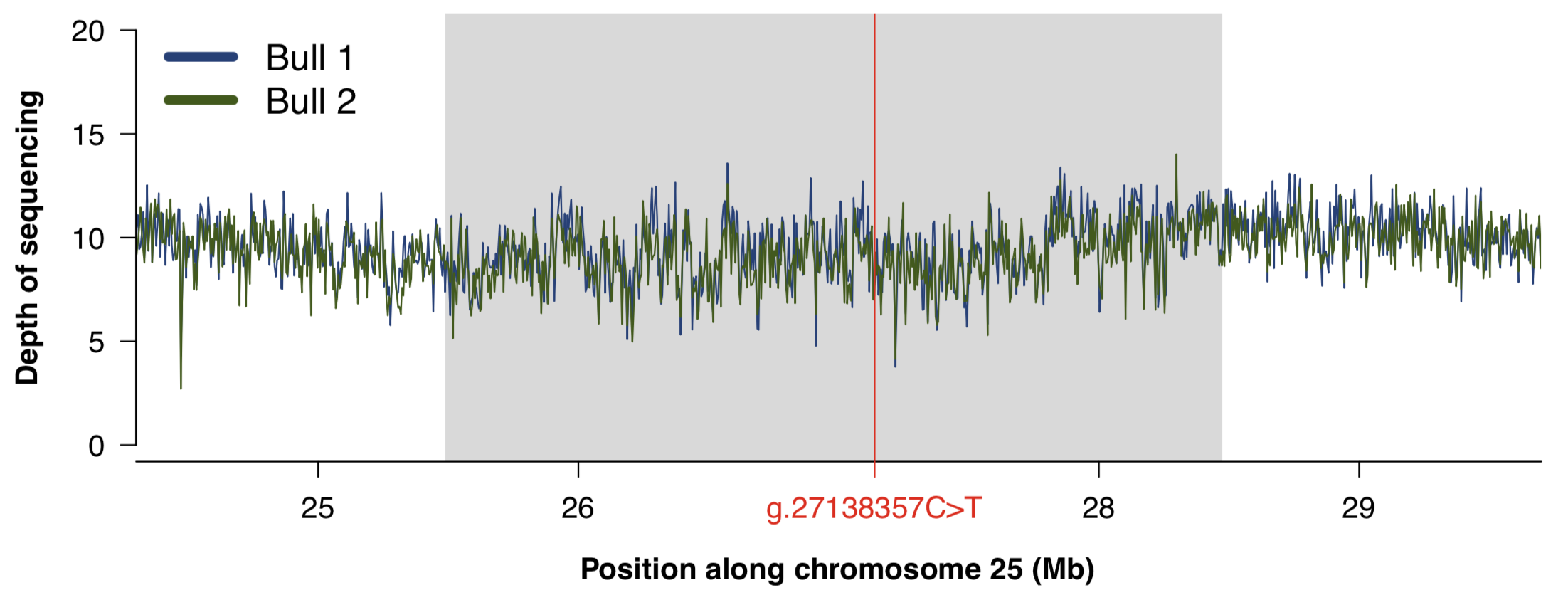

### File_S9.png

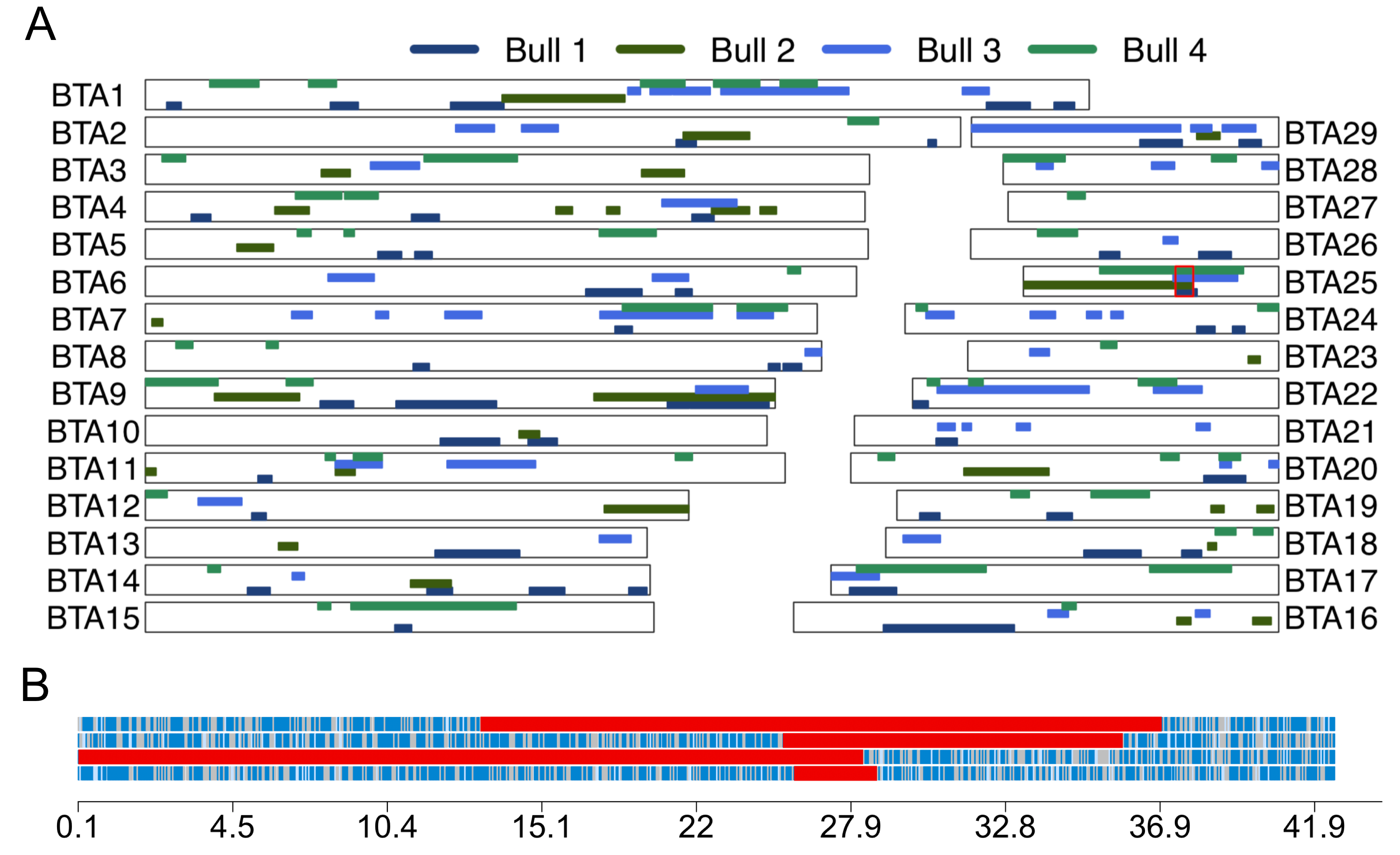
